# Preferential cross-linking of the stereospecific complex over the encounter complexes by DOPA2, a faster cross-linker than DSS

**DOI:** 10.1101/2022.06.06.494913

**Authors:** Jian-Hua Wang, Zhou Gong, Xu Dong, Shu-Qun Liu, Yu-Liang Tang, Xiaoguang Lei, Chun Tang, Meng-Qiu Dong

## Abstract

Transient protein-protein interactions are fundamental aspects of many biochemical reactions, but they are technically challenging to study. Chemical cross-linking of proteins coupled with mass spectrometry (CXMS) analysis is a powerful tool to facilitate the analysis of transient interactions. Central to this technology are chemical cross-linkers. Here, using two transient heterodimeric complexes—EIN/HPr with a *K*_D_ of 7 μM and EIIA^Glc^/EIIB^Glc^ with a *K*_D_ of 25 μM—as model systems, we compared the effects of two amine-specific homo-bifunctional cross-linkers of different cross-linking speeds. Protein cross-linking by DOPA2, a di-*ortho*-phthalaldehyde cross-linker, is 60-120 times faster than that by DSS, an N-hydroxysuccinimide ester cross-linker. We analyzed the differences in the number of cross-links identified that reflected the stereospecific complex (SC), the final lowest-energy conformational state, and that of cross-links that reflected the encounter complexes (ECs), an ensemble of short-lived intermediate conformations mediated by nonspecific electrostatic interactions. We found that the faster DOPA2 cross-linking favored the SC whereas the slower DSS cross-linking favored the ECs. We propose a mechanistic model for this intriguing observation. This study suggests that it is feasible to probe the dynamics of protein-protein interaction using cross-linkers of different cross-linking speeds.

## Introduction

Protein-protein interactions (PPIs) are essential for proper functions of proteins. The strength of a PPI can be rendered by the equilibrium dissociation constants (*K*_D_), which is equal to *k*_off_/*k*_on_, with *k*_off_ and *k*_on_ being the dissociation and association rate constant, respectively (Du *et al*. 2016). The window of biologically relevant *K*_D_ values is extremely wide, spanning 12 orders of magnitude in concentration. The PPIs are often classified as transient or stable based on their binding affinities (Perkins *et al*. 2010). Characterizing transient PPIs is technically challenging (Perkins *et al*. 2010; Qin and Gronenborn 2014). NMR is an exquisite tool for the examination of transient PPIs under physiological conditions (Vaynberg and Qin 2006; Xing *et al*. 2014; Liu *et al*. 2016), but an effective analysis of NMR spectra is often limited by the size of a protein. Other methods such as yeast two-hybrid analysis (Berggard *et al*. 2007), phage display (Smith 1985), affinity-based pull-down (Berggard *et al*. 2007), and fluorescence titration (Acuner Ozbabacan *et al*. 2011) are more successful when applied to stable interactions than to transient ones, and they do not provide direct structural information of the binding interfaces.

Chemical cross-linking of proteins coupled with mass spectrometry analysis (CXMS, XL-MS, or CLMS) has become a valuable tool for investigating the structures of protein complexes and protein-protein interactions in recent years (Yang *et al*. 2012; Liu and Heck 2015; Yu and Huang 2018; O’Reilly and Rappsilber 2018; Chavez and Bruce 2019; Wheat *et al*. 2021). In general, two residues spatially close to each other can be covalently linked by chemical cross-linkers under mild conditions. Following protease digestion, cross-linked peptide pairs can be identified by tandem mass spectrometry, which then enable localization of the cross-linked residues in protein sequences (Herzog *et al*. 2012; Wu *et al*. 2017; Zhao *et al*. 2018; Zhao *et al*. 2019; Lv *et al*. 2020). By imposing a distance restraint between cross-linked residues, which is no more than the maximal allowable distance of a cross-linker, both intra- and inter-protein interaction regions can be identified. Additionally, CXMS has the advantage of being fast and sensitive and does not require samples to be purified to homogeneity (Yu and Huang 2018; Fan *et al*. 2014).

Two interacting proteins can form a series of on-pathway or off-pathway encounter complexes (ECs) through nonspecific electrostatic interactions before they form a stereospecific complex (SC) mediated by characteristic interactions at the binding interfaces (Kozakov *et al*. 2014; Fawzi *et al*. 2010). The specific inter-protein interactions typically involve hydrogen bonding, but can also engage salt bridges, hydrophobic stacking, and *π*-stacking. It has been shown that nonspecific ECs and the SC undergo rapid interconversion (Tang *et al*. 2006; Schilder and Ubbink 2013; Anthis and Clore 2015). ECs are of short half-lives, low equilibrium population, and high heterogeneity. By forming ECs, two binding partners are kept close to each other, which accelerates the search of the complex conformational space and, hence, facilitates the formation of the SC (Tang *et al*. 2006; Dong *et al*. 2022).

SCs of large *K*_D_ values (in the μM range) and non-specific ECs are both examples of transient PPIs (La *et al*. 2013). Conventional chemical cross-linkers can capture transient PPIs. For example, studies have succeeded in using disuccinimidyl suberate (DSS) and glutaraldehyde to fix transiently formed complexes before affinity purification (Shi *et al*. 2015) or single particle cryo-electron microscopy (Kastner *et al*. 2008).

The physiochemical properties of a chemical cross-linker determine the products of cross-linking. These properties include the specificity of the reactive groups of a cross-linker (Belsom and Rappsilber 2021), the length, rigidity, and hydrophobicity/hydrophilicity of the spacer arm (Hofmann *et al*. 2015; Yu *et al*. 2020; Ding *et al*. 2016). Also important is the adopted conformation(s) of a cross-linker, which depends on whether the cross-linker is attached to a protein on one end or is completely free (Gong *et al*. 2020). Furthermore, it is suggested tentatively that the reaction kinetics of the reactive group of a cross-linker governs the capturing of a transient versus stable PPI (Belsom and Rappsilber 2021; Yang *et al*. 2018; Ziemianowicz *et al*. 2019), but this has not been investigated in a formal way.

We recently reported a class of non-hydrolyzable amine-selective di-*ortho*-phthalaldehyde (DOPA) cross-linkers, one of which is DOPA2 (Wang *et al*. 2022). The maximally possible Cα-Cα distance of DOPA2 (30.2 Å) is slightly longer than that of the widely used NHS ester cross-linker DSS (24.0 Å) (Figure 1A). Importantly, cross-linking of proteins by DOPA2 is 60-120 times faster than by DSS. We then asked whether DOPA2 might outperform DSS in capturing transient protein-protein interactions. In the current study we compared the cross-linking effects of these two cross-linkers on two transient heterodimeric complexes. Unexpectedly, we found that whether DOPA2 outperforms DSS depends on the type of transient interactions. The stereospecific complex was better depicted by the DOPA2 cross-links whereas the more heterogenous and short-lived encounter complexes were better depicted by the DSS cross-links.

**Figure 1.**
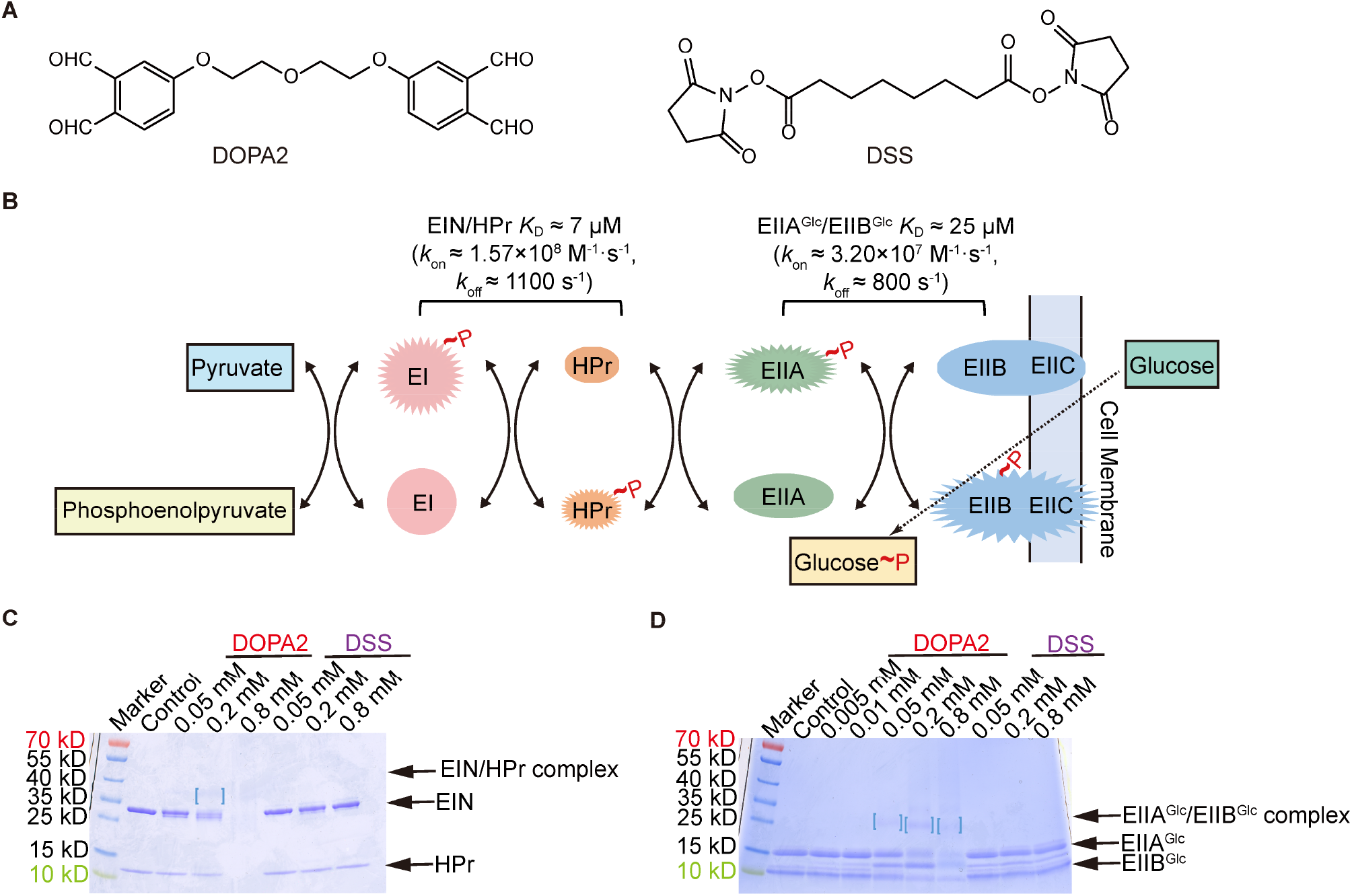
Performance of DOPA2 and DSS on transient protein complexes at the protein concentration of 1×*K*_D_. (A) The chemical structures of DOPA2 and DSS. (B) Conceptual diagram of carbohydrate transport and phosphorylation by the phosphotransferase system. (C) SDS-PAGE of DOPA2 or DSS-cross-linked EIN/HPr. The concentration of EIN/HPr is 1× *K*_D_. (D) As in C, but for the EIIA^Glc^/EIIB^Glc^ complex. Dimers were marked by square brackets. Cross-linking of EIN/HPr with 0.8 mM DOPA2 resulted in high Mw products that did not enter the separating gel.

## Results

### Transient protein complexes cross-linked by DOPA2 were more visible on SDS-PAGE gels than the DSS-linked counterparts

In the bacterial glucose phosphotransferase system (Kotrba *et al*. 2001; Deutscher *et al*. 2006), the phosphate group from phosphoenolpyruvate is transferred to glucose via Enzyme I, the phosphocarrier protein (HPr), and Enzyme II. Enzyme I is made of an N-terminal domain (EIN) and a C-terminal domain (EIC). Enzyme II (EII) comprises three functional subunits EIIA^Glc^, EIIB^Glc^, and EIIC^Glc^ (Kotrba *et al*. 2001; Deutscher *et al*. 2006). Transient interactions are responsible for the transferring of a phosphoryl group from EIN to HPr (Garrett *et al*. 1999; Garrett *et al*. 1997) and from EIIA^Glc^ to EIIB^Glc^ (Cai *et al*. 2003). The *K*_D_ values of these two protein complexes are ∼7 μM (*k*_on_ ≈ 1.57×10^8^ M^-1^ · s^-1^, *k*_off_ ≈ 1100 s^-1^) (Garrett *et al*. 1997; Suh *et al*. 2007) and ∼25 μM (*k*_on_ ≈ 3.20×10^7^ M^-1^· s^-1^, *k*_off_ ≈ 800 s^-1^) (Cai *et al*. 2003; Reizer *et al*. 1992), respectively (Figure 1B).

Consistent with the weak association between EIN and HPr and that between EIIA^Glc^ and EIIB^Glc^, DSS cross-linking yielded no discernable covalent dimer bands on the SDS-PAGE at a protein concentration of 0.25 mg/mL (for EIN/HPr) or 0.63 mg/mL (for EIIA^Glc^/EIIB^Glc^) (Figure 1C-D). Note that the protein concentrations used here are in the range for a typical cross-linking experiment. Interestingly, DOPA2 cross-linking generated a faint covalent dimer band for either complex at the same protein concentrations (Figure 1C-D). In the above experiments, the protein concentrations were set to about 1× *K*_D_, i.e., 7 µM for EIN and HPr (0.06 mg/mL and 0.19 mg/mL, respectively) and 25 µM for EIIA^Glc^ and EIIB^Glc^ (0.40 mg/mL and 0.23 mg/mL, respectively). Under these conditions, the equilibrium concentrations of the heterodimers, EIN/HPr and EIIA^Glc^/EIIB^Glc^, were 2.67 μM and 9.55 μM, respectively. In other words, only 38% of the protein subunits were in the complex form. When the protein concentrations were increased to 10× *K*_D_, the complex percentage increased to 73%, corresponding to an equilibrium concentration of 51.09 μM for the EIN/HPr heterodimer and 182.46 μM for EIIA^Glc^/EIIB^Glc^. Thus, there is a 19-fold increase in the quantity of the protein complexes relative to that at 1× *K*_D_. At 10× *K*_D_ concentration, cross-linking reactions with DOPA2 generated more prominent dimer bands on SDS-PAGE than with DSS (Figure 2A-D). Therefore, it seems that DOPA2 is better than DSS at capturing transient PPIs.

**Figure 2.**
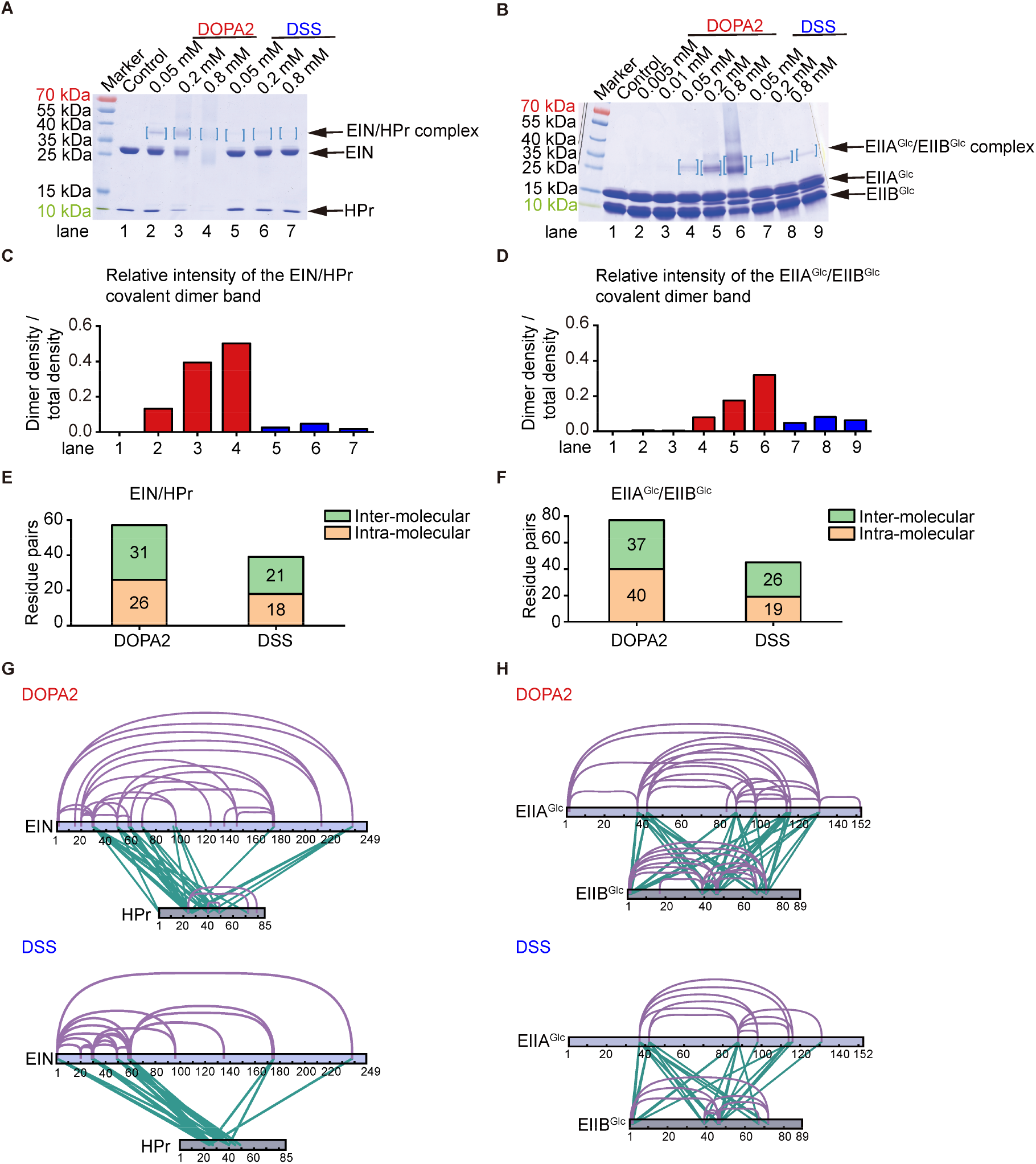
Compared to DSS, DOPA2 turned more non-covalent dimers into covalent ones that were readily visible on SDS-PAGE. (A) SDS-PAGE of DOPA2 or DSS-cross-linked EIN/HPr. The concentration of EIN/HPr is 10× *K*_D_. Dimers are marked by square brackets. (B) As in A, but for the EIIA^Glc^/EIIB^Glc^ complex. (C) The relative intensity of the EIN/HPr covalent dimer band cross-linked by DOPA2 or DSS in A. The ratio of dimer grey density to total grey density of each sample was shown in the y axis. (D) As in C, but for the EIIA^Glc^/EIIB^Glc^ covalent dimer band in B. (E) The inter-molecular or intra-molecular residue pairs identified in the EIN/HPr covalent dimer bands cross-linked by DOPA2 or DSS. (F) As in E, but for the EIIA^Glc^/EIIB^Glc^ complex. (G) DOPA2-cross-linked or DSS-cross-linked residue pairs identified from excised dimer bands that were mapped to the primary sequences of EIN/HPr complex subunits (visualized using xiNET (Combe *et al*. 2015)). (H) As in G, but for the EIIA^Glc^/EIIB^Glc^ complex. Cross-links were filtered by requiring FDR < 0.01 at the spectra level, E-value < 1×10^−3^ and spectral counts ≥ 3.

Liquid chromatography coupled with tandem mass spectrometry analysis (LC-MS/MS) of the cross-linked dimers from the gel bands indicated that the DOPA2 samples contained more intra- and inter-molecular cross-linked residue pairs (referred to as cross-links hereafter) than the DSS samples (Figure 2E-H). For example, inter-molecular cross-links between EIN and HPr numbered at 31 for DOPA2 and 21 for DSS, and those between EIIA^Glc^ and EIIB^Glc^ numbered at 37 for DOPA2 and 26 for DSS (Figure 2E-F). An analysis of the Euclidean distance of each cross-linked residue pair revealed that a higher percentage of the DOPA2 cross-links (63% for EIN/HPr, 64% for EIIA^Glc^/EIIB^Glc^) are compatible with the known complex structures than DSS cross-links (28% for EIN/HPr, 51% for EIIA^Glc^/EIIB^Glc^) (Table 1). The same conclusion holds for the solvent accessible surface distance (SASD) (Table 1).

**Table 1.**
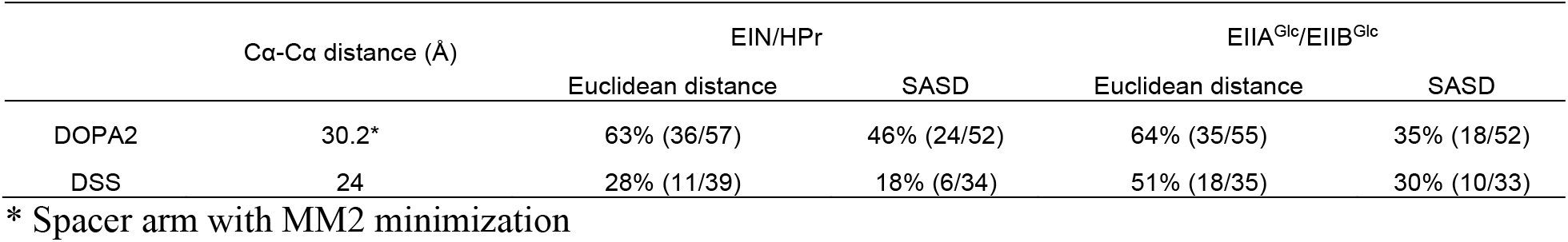
Structural compatibility rate of residue pairs (inter-molecular cross-links plus intra-molecular cross-links). SASD is short for Solvent Accessible Surface Distance. Structural models of EIN/HPr (PDB code: 3EZA) and EIIA^Glc^/EIIB^Glc^ (PDB code: 1O2F) were used as reference. Both were treated as stereospecific complexes.

### Transient protein complexes cross-linked by DSS were heterogeneous and distributed over a wide range on SDS-PAGE

In parallel, we analyzed the same cross-linking reactions without SDS-PAGE separation (Figure 3A). Interestingly, when digested in solution, the samples cross-linked with DSS yielded 23 more cross-link identifications than the samples cross-linked with DOPA2, for both intra- and inter-molecular cross-links (respectively, 40 and 13 for DSS, 24 and 6 for DOPA2) (Figure 3B). This is the opposite of the in-gel digestion result, which contained fewer DSS cross-links than DOPA2 cross-links. Furthermore, unlike the samples digested in-solution, the excised, in-gel digested covalent dimers produced more inter-than intra-molecular cross-links, either with DOPA2 or with DSS (Figure 3B). When the inter-molecular cross-links were mapped to the known structure of EIN/HPr, we found more DOPA2 cross-links than DSS cross-links bridging the interface between EIN and HPr (Figure 3C-F, denoted by orange-colored lines). Of note, the cross-links examined here all passed a stringent filter that we had applied to the pLink 2 (Chen *et al*. 2019) search results (FDR<0.01, at least four MS2 spectra at E-value < 1×10^−8^).

**Figure 3.**
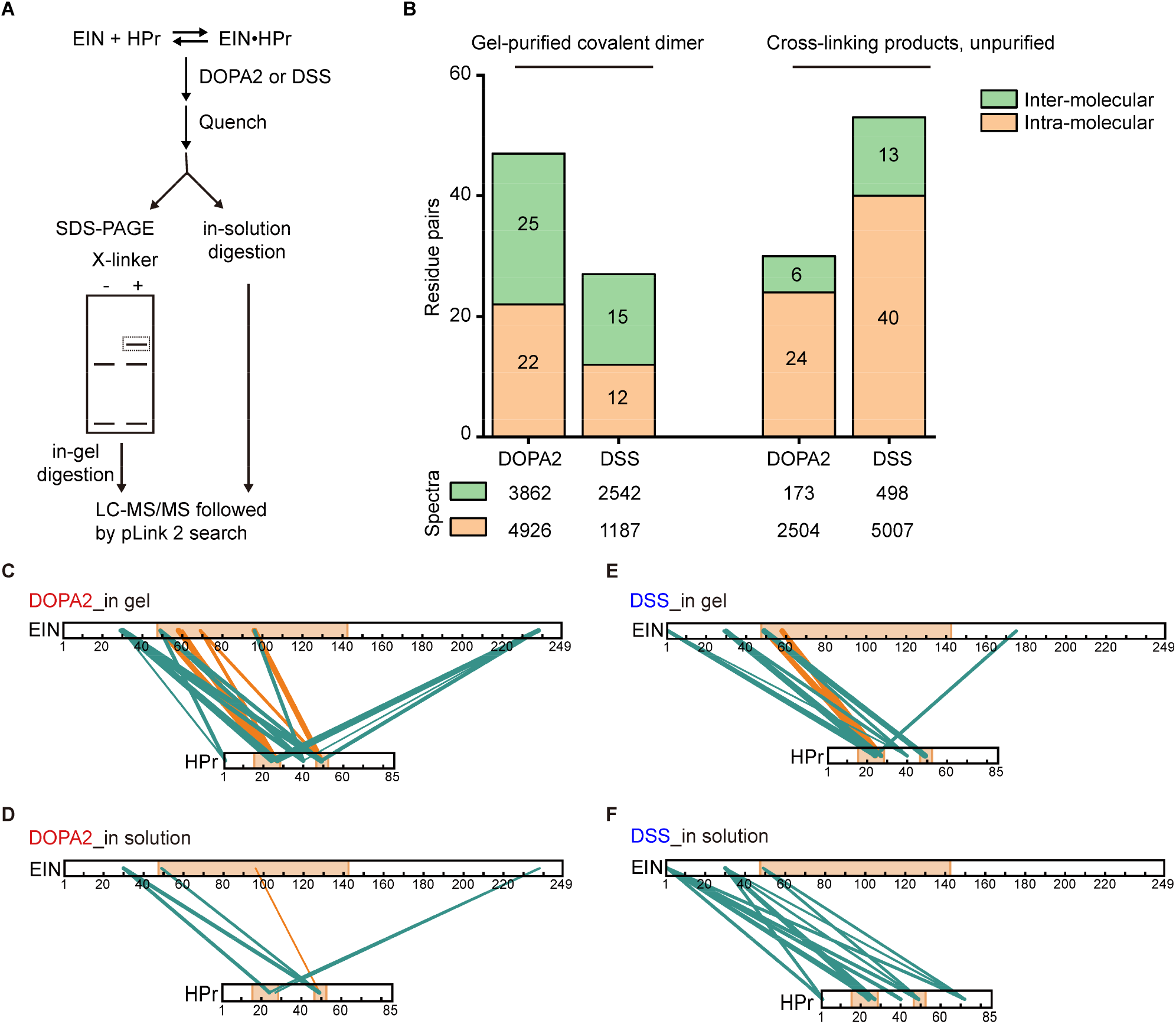
Contrasting cross-link identification results of the covalent dimer before and after isolation by SDS-PAGE. (A) Schematic diagram of in-gel digestion or in-solution digestion of EIN/HPr complex after DOPA2 or DSS cross-linking. (B) The inter-molecular or intra-molecular residue pairs identified in the EIN/HPr complex cross-linked by DOPA2 or DSS with the purification by SDS-PAGE or not. The number of spectra identified is shown below. (C and D) The inter-molecular residue pairs identified by DOPA2 in the dimer gel or in the solution were mapped on the primary sequence of EIN/HPr (visualized using xiNET (Combe *et al*. 2015)). (E and F) As in C and D, but for cross-links identified by DSS. The interactional interface of EIN and HPr on the stereospecific complex structure (PDB code: 3EZA (Garrett *et al*. 1999)) was indicated by light orange color. The green lines and the orange lines denoted the cross-links that belong to encounter complexes and stereospecific complex, respectively. The numbers of the corresponding spectra of each cross-link were indicated by the thickness of the line. Cross-links were filtered by requiring FDR < 0.01 at the spectra level, E-value < 1×10^−8^ and spectral counts > 3.

To account for the discrepancy, we systematically analyzed the cross-linking products following SDS-PAGE separation. The most prominent covalent dimer band (L3) and the gel slices above (L1-L2) and below (L4-L6) were all analyzed (Figure 4A). For intra-molecular cross-links (intra-EIN plus intra-HPr), more DSS than DOPA2 cross-links were identified (53 versus 41, respectively) from these gel slice samples (Figure 4B-C). For inter-molecular cross-links between EIN and HPr, we saw the opposite: fewer inter-molecular cross-links were identified with DSS than with DOPA2 (29 versus 33 residue pairs or 679 versus 1455 spectra, respectively) (Figure 4B-C and Figure 4D-E). Notably, inter-molecular cross-links by DOPA2 were all identified from either the visible EIN/HPr dimer band (L3) or above (L1-L2), whereas those by DSS were detected throughout the lane (L1-L5) (Figure 4B-C). The results suggest that DSS-linked dimeric complexes are more heterogeneous, spreading over a wider molecular weight range and affording a less distinct dimeric band.

**Figure 4.**
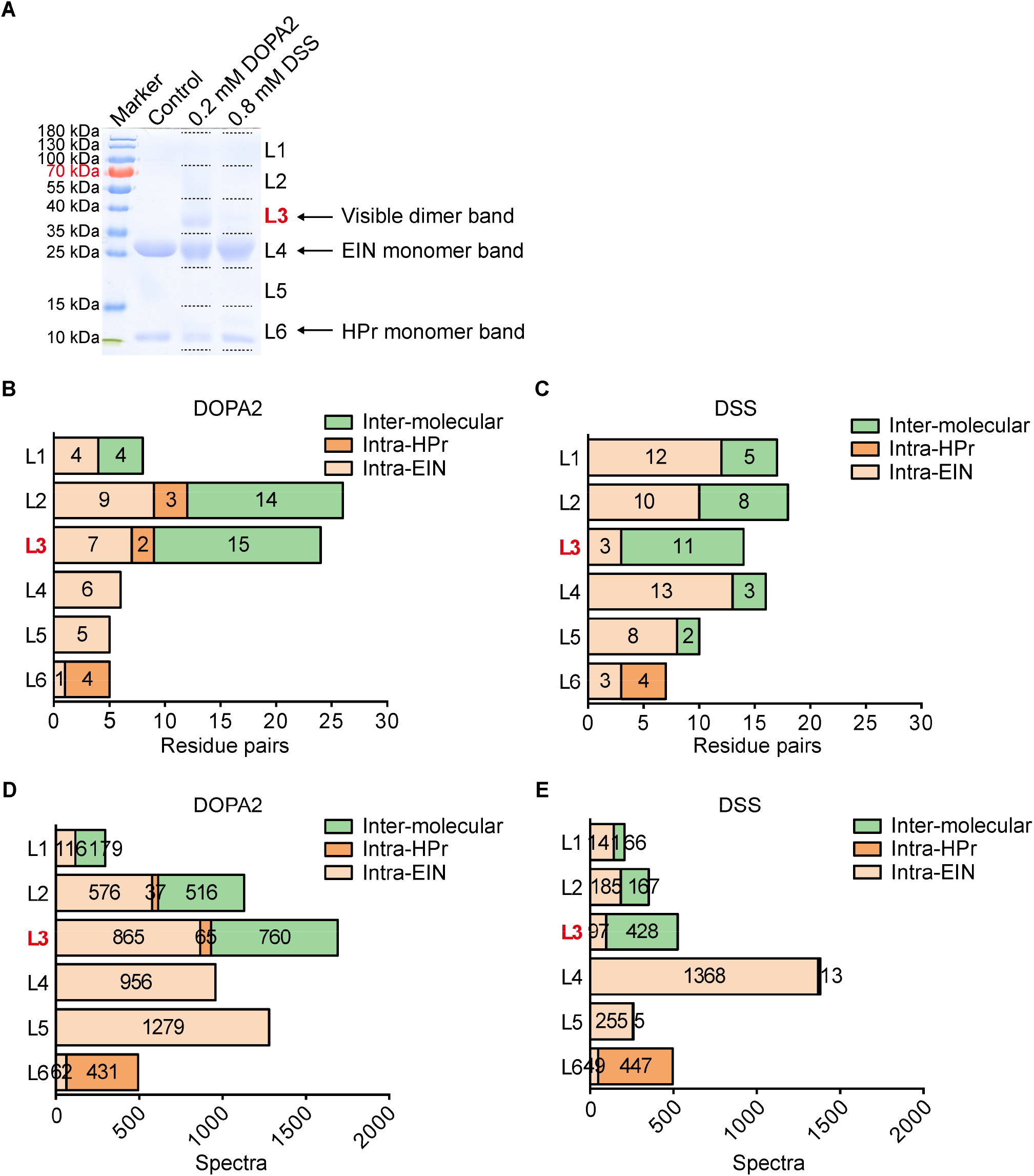
Systematic analysis of cross-linked protein species separated by SDS-PAGE. (A) SDS-PAGE of DOPA2 or DSS-cross-linked EIN/HPr and the demarcation range for systematic excision of the gel slides (L1-L6). (B and C) The number of inter- or intra-molecular residue pairs identified for DOPA2 or DSS cross-linking in L1-L6, respectively. (D and E) The number of inter- or intra-molecular spectra identified for DOPA2 or DSS cross-linking in L1-L6, respectively. Cross-links were filtered by requiring FDR < 0.01 at the spectra level, E-value < 1×10^−8^ and spectral counts in each sample ≥ 2.

### DOPA2 cross-linking favored the stereospecific complex whereas DSS cross-linking favored the encounter complexes

We showed in our previous work (Gong *et al*. 2015) that a vast majority of the EIN/HPr inter-molecular cross-links generated by BS^2^G or BS^3^, both NHS ester cross-linkers, reflected ECs. Only one out of the 13 inter-molecular BS^2^G or BS^3^ cross-links comes from SC (Gong *et al*. 2015). Fleeting ECs cannot be captured by crystallization, nor by standard NMR, but are detectable using paramagnetic NMR method (Tang *et al*. 2006). In contrast, the SC of EIN/HPr can be depicted using the standard NMR method (Tang *et al*. 2006; Garrett *et al*. 1999).

In the current study, we mapped inter-molecular cross-links on the representative structures of the ECs and the SC of EIN/HPr (Figure 5A). When the EIN/HPr samples were analyzed by LC-MS/MS without SDS-PAGE separation (in-solution samples) as done before (Gong *et al*. 2015), one of the six inter-molecular DOPA2 cross-links is consistent with the SC (17%). In contrast, none of the 13 inter-molecular DSS cross-links fitted with the SC (Figure 5B, Table 2 and Supplementary Table 1A-B). From the gel band of the covalently linked EIN/HPr dimer, four (27%) out of the 15 inter-molecular DSS cross-links represented the SC (Figure 5C, Table 2 and Supplementary Table 1C). As the SC cross-links are already favored by DOPA2, further increase by analyzing the gel band became less obvious (Figure 5C and Table 2). Nevertheless, LC-MS/MS analysis of the gel bands containing DOPA2-linked EIN/HPr showed that 8 out of 25 (32%) inter-molecular cross-links correspond to the SC (Figure 5C, Table 2 and Supplementary Table 1D). In brief, cross-links representing the SC were found enriched among the DOPA2 cross-links and among the cross-links identified from the exercised covalent dimer bands.

**Table 2.**
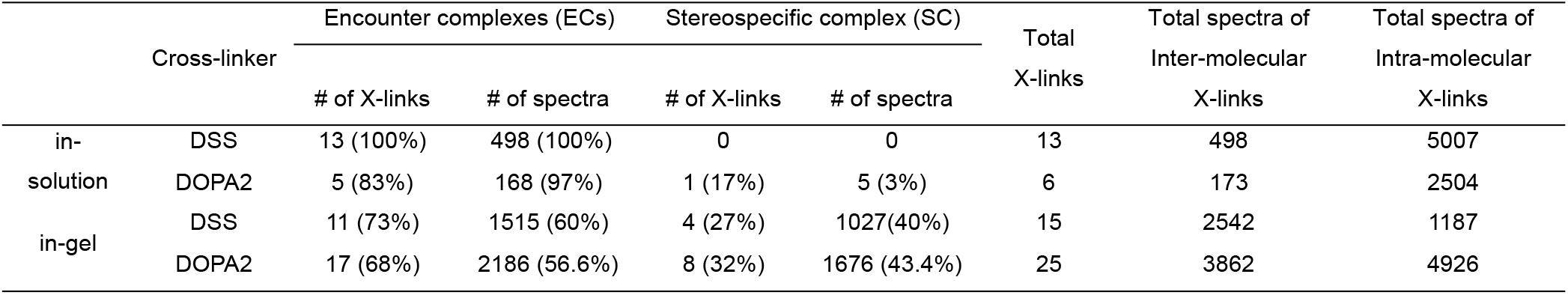
Inter-molecular cross-links identified from the EIN/HPr complex. The cross-links were filtered by requiring FDR < 0.01 at the spectra level, E-value < 1×10^−8^ and spectral counts > 3. The estimated FDR at the residue pair level is zero. The cross-links were classified according to their structural compatibility with either the stereospecific complex or the encounter complexes. The cross-linking data of 0.05, 0.2, and 0.8 mM DOPA2 or DSS were combined.

**Figure 5.**
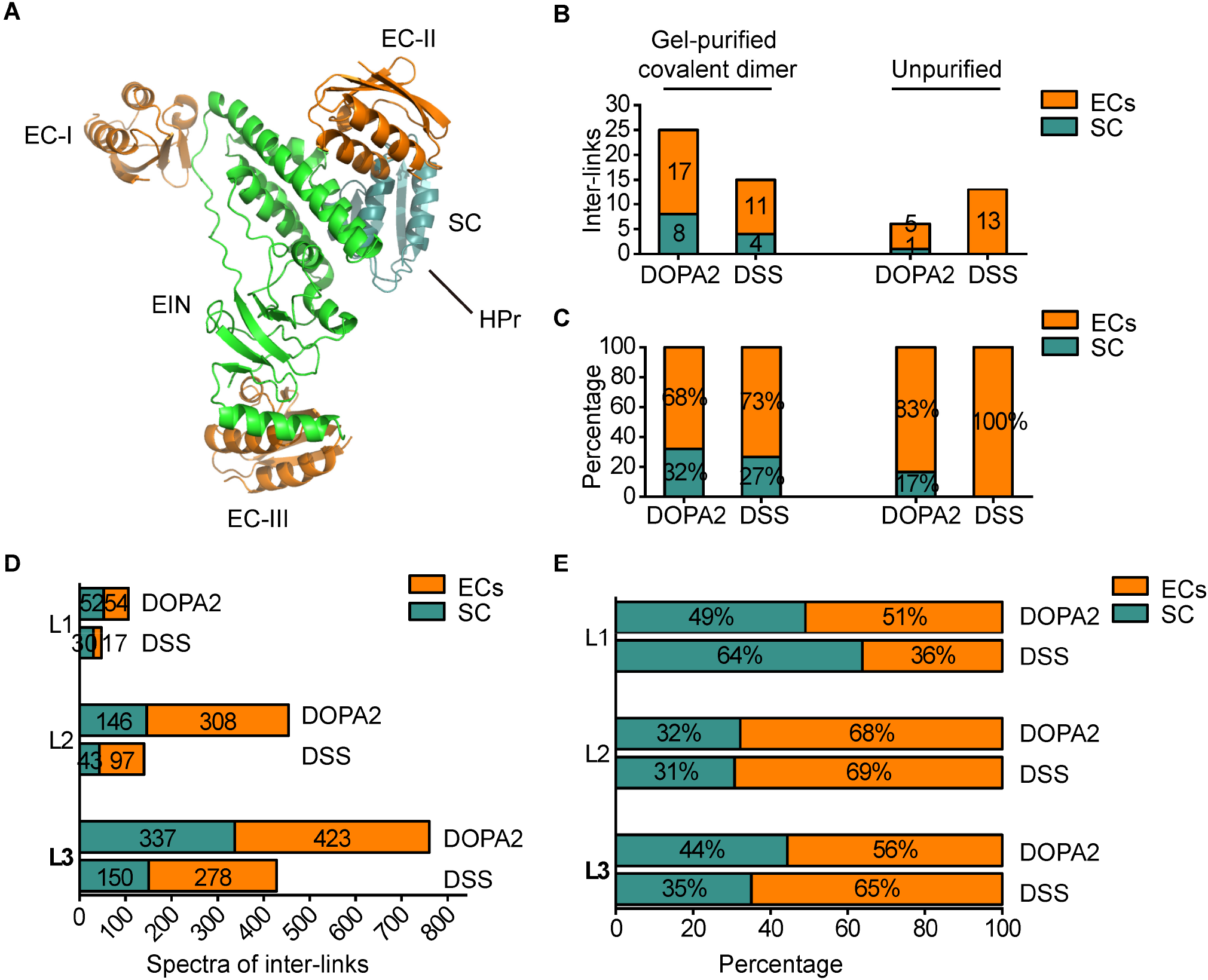
Analysis of the heterodimeric cross-links with respect to the conformations of the stereospecific and encounter complexes. (A) Three representative encounter structures (EC-I, EC-II, and EC-III) and stereospecific complex (SC, PDB code: 3EZA (Garrett *et al*. 1999) for EIN/HPr complex. (B) The number of DOPA2 or DSS cross-linked inter-links of EIN/HPr compatible with SC and ECs with purification or not. (C) The percentages of DOPA2 or DSS cross-linked inter-links of EIN/HPr compatible with SC and ECs with purification or not. (D) The number of DOPA2 or DSS cross-linked spectra of EIN/HPr compatible with SC and ECs in L1-L3, respectively. (E) The percentages of DOPA2 or DSS cross-linked spectra of EIN/HPr compatible with SC and ECs in L1-L3, respectively. Cross-links were filtered by requiring FDR < 0.01 at the spectra level, E-value < 1×10^−8^ and spectral counts in each sample ≥ 2.

We also examined the inter-molecular cross-links identified in gel slices L1-L3. Starting with the DOPA2- or DSS-linked covalent dimer bands at L3, more DOPA2 cross-links (6 residue pairs, 337 spectral counts) fitted with the SC than DSS cross-links (5 residue pairs, 150 spectral counts) (Figure 5D-E and Supplementary Table 2). This is consistent with the result shown above (Figure 5B-C) and with that from L2 (Figure 5D-E and Supplementary Table 2). The highest molecular weight region L1 had only a small number of cross-link spectra identified, diminishing but not eliminating the said difference between DOPA2 and DSS (Figure 5D and Supplementary Table 2). Together, these data suggest that DOPA2 captures the SC more readily than DSS.

### A proposed model for the differential preferences of SC and ECs by DOPA2 and DSS cross-linking

One intriguing result in this study is the differential preferences of DOPA2 and DSS towards SC and EC, respectively. Namely, for the weak EIN/HPr complex, the relatively slow cross-linking reagent DSS prefers fleeting ECs whereas the faster cross-linking reagent DOPA2 prefers the longer-lived SC. Similar to this observation, a previous study also demonstrated that N-hydroxysuccinimide esters DSS/BS^3^ trapped dynamic states in the ensemble while the fast photoactivatable diazirine-containing SDA cross-linker had better performance than DSS/BS^3^ for accurate modeling (Ziemianowicz *et al*. 2019).

The process of capturing a protein-protein interaction by cross-linking may be conceptualized as follows. We start from the intermediate state when one end of the cross-linker is already covalently attached to either protein (A or B) in a transient complex A•B. For the subsequent inter-molecular cross-linking reaction, the likelihood for the formation of an inter-molecular cross-link will be determined by the reaction rate of the amine-targeting group as well as the half-life of the A•B complex. If the reaction rate is not fast enough compared to the half-life of the A•B complex, no inter-molecular cross-link will form in this round of association. However, this “planted” cross-linker still has a chance to form an inter-molecular cross-link in the next round of association before it loses reactivity due to the attack of an intra-molecular amine group or a quenching reagent.

As shown in Figure 6, we propose that DSS or BS^3^ cross-linking reactions are too slow to capture either the SC or the fleeting ECs in one association/dissociation cycle. As a result, the observed cross-links may come from subsequent association events. As the SC and the ECs undergo rapid interconversion, it is not surprising that a DSS or BS^3^ mono-linked protein can be attached to its partner protein in conformations other than the SC. This is in line with an earlier work demonstrating that a slow reacting sulfonyl fluoride group can capture weak and transient interactions once planted onto a protein (Yang *et al*. 2018). This explains the observation that the number of inter-molecular DSS or BS^3^ cross-links identified from EIN/HPr is small and most of them represent ECs. In contrast, DOPA2 has a faster reaction rate and therefore can be more efficient in capturing the SC before the two subunits dissociate. Indeed, our recent NMR analysis showed that the EIN/HPr complex has a *k*_off_ of 8900 s^-1^, corresponding to a lifetime of ∼100 µs (Dong *et al*. 2022). As ECs exist as an ensemble of conformations while the SC represents the final lowest-energy conformational state, cross-linked ECs are more heterogeneous than the cross-linked SC, and therefore are more spread-out and less visible on SDS-PAGE.

**Figure 6.**
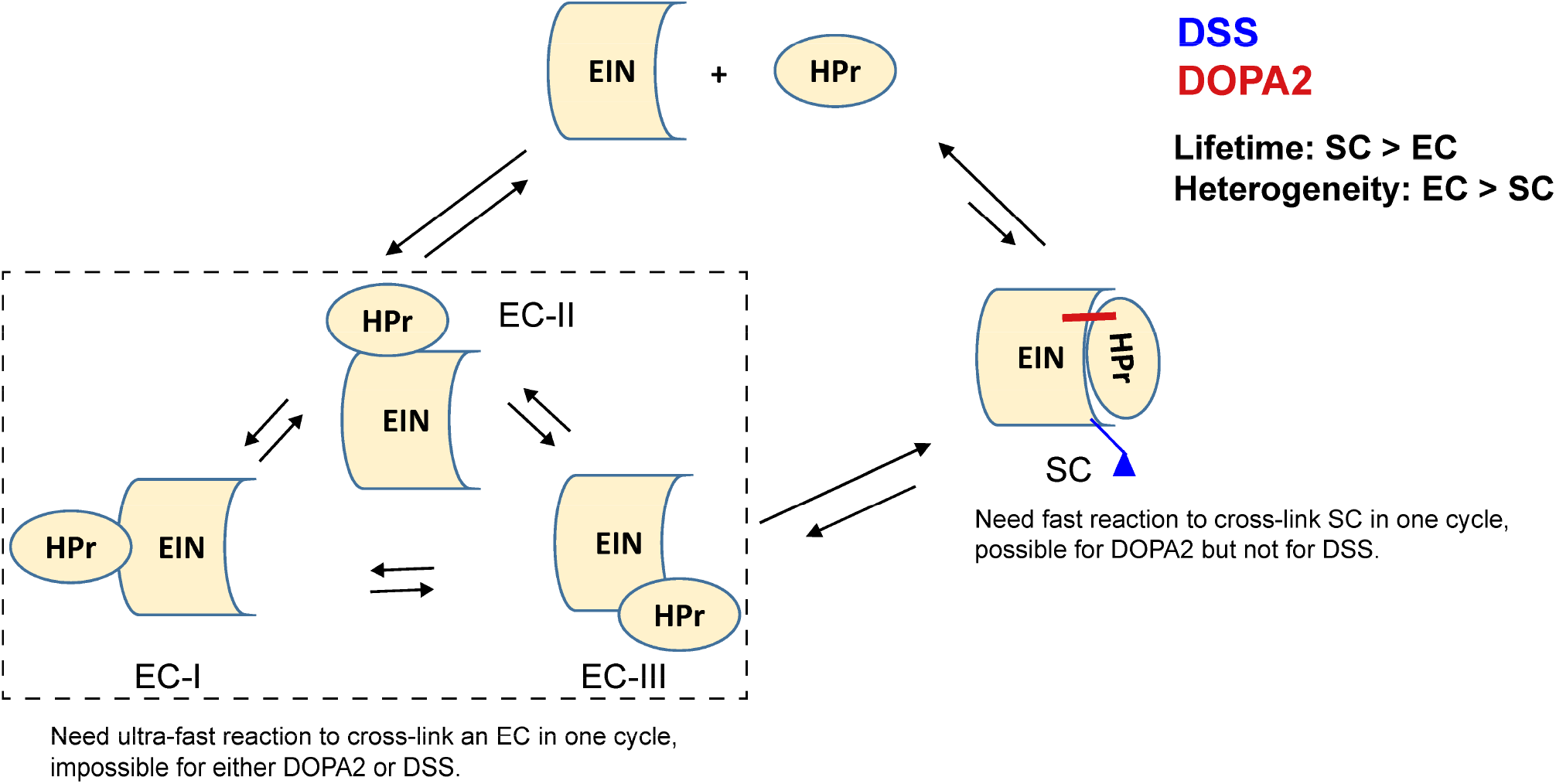
A proposed model for the differential preferences of SC and ECs by DOPA2 and DSS cross-linking. (1) An EC is too transient for either DOPA2 or DSS to capture immediately. (2)Compared to DSS, the fast-reacting cross-linker DOPA2 has a higher chance of capturing the stereospecific complex of EIN/HPr, by forming an inter-molecular cross-link before the two subunits dissociate. (3)Cross-linking by DSS is too slow to capture an EIN/HPr complex in one shot. DSS cross-linking probably takes place in two steps: first a DSS mono-link forms, then in one or more dissociation/association cycles, the mono-link planted on a subunit turns into a cross-link. Between the mono-link and the cross-link, the protein has many opportunities to sample different conformational states, including ECs.

## Discussion

In this study, we have demonstrated that under the typical, near-physiological conditions, DOPA2 is advantageous over DSS in capturing transiently interacting proteins for the structural characterization of a stereospecific complex. The idea that a faster cross-linker is advantageous in capturing a transient PPI is likely correct, but the truth is much more nuanced than expected. “Fast” and “slow” are relative terms. DOPA2 is perhaps fast enough for the SC of EIN/HPr, but certainly not for the fleeting ECs. Although too slow for either the SC or ECs, DSS counterintuitively has a better ability to capture ECs, probably by going through multiple cycles of association and dissociation. This may be a unique advantage of DSS, but as cross-linking modifies protein surface by conjugating a small molecule to lysine side chains, which could modify how proteins interact with one another, further cross-validating experiments are warranted in assessing the correctness of the captured ECs.

Since the structures of ECs of EIIA^Glc^/EIIB^Glc^ are not yet available, we cannot validate whether cross-links originated from the ECs were enriched in the DSS cross-links of this heterodimer, but we think it is a reasonable hypothesis because the contrasting behaviors of DSS and DOPA2 on EIN/HPr were also seen on EIIA^Glc^/EIIB^Glc^. For either heterodimeric complex, more DOPA2 cross-links (>60%) fitted with the structure of the SC than did the DSS cross-links (28-51%) (Table 1). Further, DOPA2 cross-linking generated a covalent dimer band that is much more distinct on SDS-PAGE than did DSS cross-linking for both EIN/HPr and EIIA^Glc^/EIIB^Glc^ (Figure 1 and Figure 2). For EIN/HPr, the SC is highly represented in this dimer band (Table 2) as discussed above.

We have noticed in our previous analysis of six model proteins (aldolase, BSA, catalase, GST, lysozyme, and myosin), the DOPA2 cross-links have a higher compatibility rate with the determined structures of these proteins than the DSS cross-links, as do the cross-links of DOPA-C_2_, another DOPA cross-linker (Wang *et al*. 2022). It is unlikely that this difference is due to a longer cross-linking distance of DOPA2 (30.2Å), because DOPA-C_2_ (24.9 Å) is close to DSS (24.0 Å) in this regard. Based on the findings from the current study and that the determined structures are biased towards the lowest-energy conformational states (e.g. the SC), we propose that the difference in the structural compatibility rate may be accounted for, to some extent, by the faster cross-linking speed of DOPA cross-linkers relative to the NHS ester cross-linker DSS.

## Experimental Section

### Materials and reagents

DSS, tris(2-carboxyethyl) phosphine (TCEP), Dithiothreitol (DTT), and 2-Iodoacetamide (IAA) were purchased from Pierce biotechnology (Thermo Scientific). Dimethylsulfoxide (DMSO), HEPES, NaCl, urea, CaCl_2,_ and methylamine were purchased from Sigma-Aldrich. Acetonitrile (ACN), formic acid (FA), acetone, and ammonium bicarbonate were purchased from J.T. Baker. Trypsin (gold mass spectrometry grade) was purchased from Promega. DOPA2 was synthesized by Professor Lei Xiaoguang Laboratory of Peking University, China.

### Preparation of protein samples

The N-terminal domain of *E. coli* enzyme I (EIN, residues 1-249) and the histidine-containing phosphorcarrier protein HPr, EIIA^Glc^ and EIIB^Glc^ were purified as previously described (Xing *et al*. 2014; Garrett *et al*. 1999). Eluted proteins were exchanged into 20 mM HEPES, 150 mM NaCl, pH 7.4.

### Protein cross-linking

The *K*_D_ value of EIN/HPr complex was ∼7 μM (Suh *et al*. 2007). EIN/HPr complexes were diluted to 1× *K*_D_ (0.25 mg/mL) and 10× *K*_D_ (2.5 mg/mL), and then were cross-linked with DOPA2 at the final concentration of 0.05 mM, 0.2 mM, 0.8 mM at room temperature for 10 minutes, respectively. EIN/HPr complexes were also cross-linked with DSS at the final concentration of 0.05 mM, 0.2 mM, 0.8 mM at room temperature for one hour.

The *K*_D_ value of EIIA^Glc^/EIIB^Glc^ complex was estimated to be ∼25 μM. According to a previous study, EIIA^Glc^/EIIB^Glc^ has a *K*m value of 1.7-25 μM (Reizer *et al*. 1992). We did not succeed in obtaining an accurate *K*_D_ value by either surface plasmon resonance assay or ELISA. However, it is clear that the binding is weak, and the *K*_D_ value is likely greater than 10 μM. EIIA^Glc^/EIIB^Glc^ complexes were diluted to 1× *K*_D_ (0.63 mg/mL) and 10× *K*_D_ (6.3 mg/mL), and then were cross-linked with DOPA2 at the final concentration of 0.005 mM, 0.01 mM, 0.05 mM, 0.2 mM, 0.8 mM at room temperature for 10 minutes, respectively. EIIA^Glc^/EIIB^Glc^ complexes were also cross-linked with DSS at the final concentration of 0.05 mM, 0.2 mM, 0.8 mM at room temperature for one hour.

### In-solution digestion

Cross-linked proteins were precipitated with 4-fold volume of acetone for at least 30 minutes at −20 °C. The pellets were air dried and then dissolved, assisted by sonication, in 8 M urea, 20 mM methylamine, 100 mM Tris, pH 8.5. After reduction (5 mM TCEP, RT, 20 min) and alkylation (10 mM IAA, RT, 15 min in the dark), the samples were diluted to 2 M urea with 100 mM Tris, pH 8.5. Denatured proteins were digested by trypsin at a 1/50 (w/w) enzyme/substrate ratio at 37 °C for 16-18 h, and the reactions were quenched with 5% formic acid (final conc.).

### In-gel digestion

The target bands in the one-dimensional glycine gel were excised manually from the gel slab and cut into pieces. Briefly, after fully washed with destaining solution and ddH_2_O, the gel pieces were in-gel reduced (10 mM DTT, 56 °C, 40 min) and alkylated (55 mM IAA, RT, 60 min in the dark) and then dehydrated by 100% ACN. Gel pieces were further rehydrated (50 mM NH_4_HCO_3_, 10 ng/μl trypsin) and digested for 16-18 h. The peptides were twice extracted from the gel by extraction solution I (50% ACN, 5% FA) and extraction solution II (75% ACN, 5% FA), respectively. The extracted digests were combined and the sample volume was reduced to about 10 μl in speedvac for MS analysis.

### LC-MS analysis

All proteolytic digestions of proteins were analyzed using an EASY-nLC 1000 system (Thermo Fisher Scientific) interfaced with an HF Q-Exactive mass spectrometer (Thermo Fisher Scientific). Peptides were loaded on a pre-column (75 μm ID, 4 cm long, packed with ODS-AQ 12 nm-10 μm beads) and separated on an analytical column (75 μm ID, 12 cm long, packed with Luna C18 1.9 μm 100 Å resin). EIN/HPr and EIIA^Glc^/EIIB^Glc^ complexes were injected and separated with a 75 min linear gradient at a flow rate of 200 nl/min as follows: 0-5% B in 1 min, 5-35% B in 59 min, 35-100% B in 5 min, 100% B for 10 min (A = 0.1% FA, B = 100% ACN, 0.1% FA). The top fifteen most intense precursor ions from each full scan (resolution 60,000) were isolated for HCD MS2 (resolution 15,000; NCE 27) with a dynamic exclusion time of 30 s. Precursors with 1+, 2+, more than 6+, or unassigned charge states were excluded.

### Identification of cross-links with pLink 2

The search parameters used for pLink 2 (Chen *et al*. 2019) were as follows: instrument, HCD; precursor mass tolerance, 20 ppm; fragment mass tolerance 20 ppm; cross-linker DOPA2 (cross-linking sites K and protein N-terminus, cross-link mass-shift 334.084, mono-link w/o hydrazine mass-shift 352.096, mono-link w/t hydrazine mass-shift 348.111), cross-linker DSS (cross-linking sites K and protein N-terminus, cross-link mass-shift 138.068, mono-link mass-shift 156.079); fixed modification Carbamidomethyl[C]; variable modifications Deamidated[N], Deamidated[Q], and Oxidation[M]; peptide length, minimum 6 amino acids and maximum 60 amino acids per chain; peptide mass, minimum 600 and maximum 6,000 Da per chain; enzyme, trypsin, with up to three missed cleavage sites per cross-link. Protein sequences of model proteins were used for database searching. The results were filtered by requiring a spectral false identification rate < 0.01.

### Cα-Cα distance calculations

The Cα-Cα Euclidean distances were measured using PyMOL in a PDB file. The PDB files we use are as follows: EIN/HPr (3EZA (Garrett *et al*. 1999)), EIIA^Glc^/EIIB^Glc^ (1O2F (Cai *et al*. 2003)). The Cα-Cα Solvent Accessible Surface Distance (SASD) were calculated using Jwalk (Matthew Allen Bullock *et al*. 2016). If the SASD of cross-linked residue pairs cannot be calculated due to a lack of surface accessibility, these residue pairs are excluded from calculation. When calculating structural compatibility, the distance cut-offs are 30.2 Å for DOPA2, and 24.0 Å for DSS.

## Acknowledgment

We thank Yulu Li and Dr. Jianhua Sui for the help in determining the *K*_D_ value of EIIA^Glc^/EIIB^Glc^. We gratefully acknowledge financial support from the following grants: Development of Major Scientific Instruments and Equipment of China (2020YFF01014505 to M.-Q.D.) and the National Key Research and Development Program (2018YFA0507700 to C.T.).

## Supplementary Information

**Supplementary Table 1.** The inter-molecular cross-links identified by DOPA2 or DSS in the EIN/HPr complex. (A) The inter-molecular cross-links identified by DSS in the solution. (B) As in A, but for cross-links identified by DOPA2. (C) The inter-molecular cross-links identified by DSS in the gel. (D) As in C, but for cross-links identified by DOPA2. Cross-links were filtered by requiring FDR < 0.01 at the spectra level, E-value < 1×10^−8^ and spectral counts > 3. The estimated FDR at the residue pair level is zero.

| Linked aa positions | Total Spec | Best E-value | Ca-Ca distance (Å) (PDB code: 3EZA) | label |
| --- | --- | --- | --- | --- |
| EIN(49)-HPr(49) | 22 | 1.78E-20 | 26.48 | in-solution |
| EIN(30)-HPr(49) | 13 | 3.73E-37 | 29.90 | in-solution |
| EIN(49)-HPr(72) | 20 | 4.17E-16 | 35.22 | in-solution |
| HPr(27)-EIN(30) | 27 | 2.66E-19 | 35.30 | in-solution |
| HPr(24)-EIN(30) | 119 | 2.86E-17 | 36.78 | in-solution |
| EIN(1)-HPr(49) | 48 | 4.34E-32 | 44.96 | in-solution |
| HPr(1)-EIN(30) | 32 | 2.72E-19 | 46.92 | in-solution |
| EIN(30)-HPr(72) | 9 | 3.71E-21 | 47.21 | in-solution |
| EIN(1)-HPr(40) | 58 | 3.18E-11 | 48.40 | in-solution |
| EIN(1)-HPr(24) | 18 | 8.06E-17 | 49.07 | in-solution |
| EIN(1)-HPr(27) | 63 | 5.71E-22 | 53.15 | in-solution |
| EIN(1)-HPr(72) | 32 | 1.17E-27 | 60.88 | in-solution |
| EIN(1)-HPr(1) | 37 | 1.26E-31 | 64.70 | in-solution |

B
| Linked aa positions | Total Spec | Best E-value | Ca-Ca distance (Å) (PDB code: 3EZA) | label |
| --- | --- | --- | --- | --- |
| HPr(49)-EIN(96) | 5 | 1.92E-19 | 24.33 | in-solution |
| EIN(49)-HPr(49) | 21 | 1.05E-17 | 26.48 | in-solution |
| EIN(30)-HPr(49) | 83 | 3.73E-31 | 29.90 | in-solution |
| HPr(24)-EIN(30) | 47 | 1.95E-12 | 36.78 | in-solution |
| HPr(24)-EIN(238) | 11 | 5.31E-16 | 56.10 | in-solution |
| HPr(27)-EIN(238) | 6 | 1.66E-20 | 61.06 | in-solution |

C
| Linked aa positions | Total Spec | Best E-value | Ca-Ca distance (Å)<br>(PDB code: 3EZA) | label |
| --- | --- | --- | --- | --- |
| HPr(24)-EIN(58) | 526 | 3.21E-36 | 15.38 | in-gel |
| HPr(27)-EIN(58) | 238 | 5.11E-14 | 19.62 | in-gel |
| HPr(24)-EIN(49) | 94 | 1.58E-12 | 22.32 | in-gel |
| HPr(27)-EIN(49) | 169 | 4.16E-31 | 23.42 | in-gel |
| EIN(49)-HPr(49) | 702 | 2.37E-24 | 26.48 | in-gel |
| HPr(24)-EIN(175) | 43 | 2.83E-22 | 34.05 | in-gel |
| HPr(27)-EIN(30) | 156 | 2.81E-16 | 35.30 | in-gel |
| HPr(40)-EIN(49) | 27 | 4.61E-24 | 36.39 | in-gel |
| HPr(24)-EIN(30) | 303 | 5.70E-20 | 36.78 | in-gel |
| HPr(27)-EIN(29) | 36 | 2.84E-26 | 36.85 | in-gel |
| HPr(24)-EIN(29) | 56 | 2.87E-09 | 37.81 | in-gel |
| EIN(30)-HPr(40) | 44 | 4.52E-21 | 38.01 | in-gel |
| EIN(1)-HPr(40) | 5 | 1.94E-10 | 48.40 | in-gel |
| EIN(1)-HPr(24) | 81 | 4.86E-13 | 49.07 | in-gel |
| EIN(1)-HPr(27) | 62 | 3.69E-21 | 53.15 | in-gel |

D
| Linked aa positions | Total Spec | Best E-value | Ca-Ca distance (Å)<br>(PDB code: 3EZA) | label |
| --- | --- | --- | --- | --- |
| HPr(24)-EIN(58) | 233 | 8.11E-40 | 15.38 | in-gel |
| HPr(27)-EIN(58) | 504 | 4.98E-23 | 19.62 | in-gel |
| HPr(24)-EIN(60) | 15 | 1.31E-09 | 20.21 | in-gel |
| HPr(27)-EIN(69) | 45 | 1.65E-14 | 20.35 | in-gel |
| HPr(49)-EIN(69) | 28 | 1.58E-18 | 22.20 | in-gel |
| HPr(24)-EIN(49) | 127 | 2.89E-33 | 22.32 | in-gel |
| HPr(27)-EIN(49) | 86 | 3.44E-23 | 23.42 | in-gel |
| HPr(49)-EIN(96) | 638 | 3.05E-41 | 24.33 | in-gel |
| EIN(49)-HPr(49) | 78 | 6.50E-18 | 26.48 | in-gel |
| EIN(30)-HPr(49) | 623 | 1.18E-34 | 29.90 | in-gel |
| EIN(29)-HPr(49) | 24 | 6.25E-16 | 30.92 | in-gel |
| HPr(40)-EIN(96) | 56 | 1.95E-10 | 31.79 | in-gel |
| HPr(27)-EIN(30) | 274 | 2.13E-21 | 35.30 | in-gel |
| HPr(40)-EIN(49) | 77 | 3.25E-28 | 36.39 | in-gel |
| HPr(24)-EIN(30) | 209 | 8.37E-23 | 36.78 | in-gel |
| HPr(27)-EIN(29) | 78 | 3.22E-26 | 36.85 | in-gel |
| HPr(24)-EIN(29) | 37 | 1.21E-10 | 37.81 | in-gel |
| EIN(30)-HPr(40) | 130 | 2.27E-31 | 38.01 | in-gel |
| EIN(29)-HPr(40) | 24 | 5.62E-16 | 39.11 | in-gel |
| HPr(1)-EIN(49) | 66 | 6.96E-26 | 41.09 | in-gel |
| HPr(1)-EIN(30) | 7 | 2.05E-19 | 46.92 | in-gel |
| HPr(24)-EIN(238) | 229 | 2.10E-13 | 56.10 | in-gel |
| HPr(49)-EIN(238) | 65 | 1.76E-20 | 56.45 | in-gel |
| HPr(40)-EIN(238) | 4 | 3.15E-09 | 60.32 | in-gel |
| HPr(27)-EIN(238) | 205 | 2.49E-32 | 61.06 | in-gel |

**Supplementary Table 2.** Inter-molecular cross-links between EIN and HPr identified from gel slices L1-L3. The cross-links were filtered by requiring FDR < 0.01 at the spectra level, E-value < 1×10^−8^ and spectral counts in each sample ≥ 2. The estimated FDR at the residue pair level is zero. The cross-links were classified according to their structural compatibility with either the stereospecific complex or the encounter complexes.

| Cross-linker | Gel slices | Encounter complexes (ECs) |  | Stereospecific complex (SC) |  |
| --- | --- | --- | --- | --- | --- |
|  |  | # of X-links | # of spectra | # of X-links | # of spectra |
| DOPA2 | L1 | 1 | 54 | 2 | 52 |
| DOPA2 | L2 | 8 | 308 | 5 | 146 |
| DOPA2 | L3 | 9 | 423 | 6 | 337 |
| DSS | L1 | 1 | 17 | 3 | 30 |
| DSS | L2 | 5 | 97 | 2 | 43 |
| DSS | L3 | 6 | 278 | 5 | 150 |

## References

[1] Du, X, Li, Y, Xia, YL, Ai, SM, Liang, J, Sang, P, Ji, XL, Liu, SQ (2016) Insights into protein-ligand interactions: mechanisms, models, and methods. Int J Mol Sci 17.

[2] Perkins, JR, Diboun, I, Dessailly, BH, Lees, JG, Orengo, C (2010) Transient protein-protein interactions: structural, functional, and network properties. Structure 18: 1233–1243.

[3] Qin, J, Gronenborn, AM (2014) Weak protein complexes: challenging to study but essential for life. FEBS J 281: 1948–1949.

[4] Vaynberg, J, Qin, J (2006) Weak protein-protein interactions as probed by NMR spectroscopy. Trends Biotechnol 24: 22–27.

[5] Xing, Q, Huang, P, Yang, J, Sun, JQ, Gong, Z, Dong, X, Guo, DC, Chen, SM, Yang, YH, Wang, Y, Yang, MH, Yi, M, Ding, YM, Liu, ML, Zhang, WP, Tang, C (2014) Visualizing an ultra-weak protein-protein interaction in phosphorylation signaling. Angew Chem Int Ed Engl 53: 11501–11505.

[6] Liu, Z, Gong, Z, Dong, X, Tang, C (2016) Transient protein-protein interactions visualized by solution NMR. Biochim Biophys Acta 1864: 115–122.

[7] Berggard, T, Linse, S, James, P (2007) Methods for the detection and analysis of protein-protein interactions. Proteomics 7: 2833–2842.

[8] Smith, GP (1985) Filamentous fusion phage: novel expression vectors that display cloned antigens on the virion surface. Science 228: 1315–1317.

[9] Acuner Ozbabacan, SE, Engin, HB, Gursoy, A, Keskin, O (2011) Transient protein-protein interactions. Protein Eng Des Sel 24: 635–648.

[10] Yang, B, Wu, YJ, Zhu, M, Fan, SB, Lin, J, Zhang, K, Li, S, Chi, H, Li, YX, Chen, HF, Luo, SK, Ding, YH, Wang, LH, Hao, Z, Xiu, LY, Chen, S, Ye, K, He, SM, Dong, MQ (2012) Identification of cross-linked peptides from complex samples. Nat Methods 9: 904–906.

[11] Liu, F, Heck, AJ (2015) Interrogating the architecture of protein assemblies and protein interaction networks by cross-linking mass spectrometry. Curr Opin Struct Biol 35: 100–108.

[12] Yu, C, Huang, L (2018) Cross-linking mass spectrometry: an emerging technology for interactomics and structural biology. Anal Chem 90: 144–165.

[13] O’Reilly, FJ, Rappsilber, J (2018) Cross-linking mass spectrometry: methods and applications in structural, molecular and systems biology. Nat Struct Mol Biol 25: 1000–1008.

[14] Chavez, JD, Bruce, JE (2019) Chemical cross-linking with mass spectrometry: a tool for systems structural biology. Curr Opin Chem Biol 48: 8–18.

[15] Wheat, A, Yu, C, Wang, X, Burke, AM, Chemmama, IE, Kaake, RM, Baker, P, Rychnovsky, SD, Yang, J, Huang, L (2021) Protein interaction landscapes revealed by advanced in vivo cross-linking-mass spectrometry. Proc Natl Acad Sci U S A 118: e2023360118.

[16] Herzog, F, Kahraman, A, Boehringer, D, Mak, R, Bracher, A, Walzthoeni, T, Leitner, A, Beck, M, Hartl, FU, Ban, N, Malmstrom, L, Aebersold, R (2012) Structural probing of a protein phosphatase 2A network by chemical cross-linking and mass spectrometry. Science 337: 1348–1352.

[17] Wu, S, Tan, D, Woolford, JL, Jr., Dong, MQ, Gao, N (2017) Atomic modeling of the ITS2 ribosome assembly subcomplex from cryo-EM together with mass spectrometry-identified protein-protein crosslinks. Protein Sci 26: 103–112.

[18] Zhao, Q, Zhou, H, Chi, S, Wang, Y, Wang, J, Geng, J, Wu, K, Liu, W, Zhang, T, Dong, MQ, Wang, J, Li, X, Xiao, B (2018) Structure and mechanogating mechanism of the Piezo1 channel. Nature 554: 487–492.

[19] Zhao, K, Cheng, S, Miao, N, Xu, P, Lu, X, Zhang, Y, Wang, M, Ouyang, X, Yuan, X, Liu, W, Lu, X, Zhou, P, Gu, J, Zhang, Y, Qiu, D, Jin, Z, Su, C, Peng, C, Wang, JH, Dong, MQ, Wan, Y, Ma, J, Cheng, H, Huang, Y, Yu, Y (2019) A Pandas complex adapted for piRNA-guided transcriptional silencing and heterochromatin formation. Nat Cell Biol 21: 1261–1272.

[20] Lv, L, Chen, P, Cao, L, Li, Y, Zeng, Z, Cui, Y, Wu, Q, Li, J, Wang, JH, Dong, MQ, Qi, X, Han, T (2020) Discovery of a molecular glue promoting CDK12-DDB1 interaction to trigger cyclin K degradation. Elife 9: e59994.

[21] Fan, SB, Wu, YJ, Yang, B, Chi, H, Meng, JM, Lu, S, Zhang, K, Wu, L, Sun, RX, Dong, MQ, He, SM (2014) A new approach to protein structure and interaction research: chemical cross-linking in combination with mass spectrometry. Progress in Biochemistry and Biophysics 41: 1109–1125.

[22] Kozakov, D, Li, K, Hall, DR, Beglov, D, Zheng, J, Vakili, P, Schueler-Furman, O, Paschalidis, I, Clore, GM, Vajda, S (2014) Encounter complexes and dimensionality reduction in protein-protein association. Elife 3: e01370.

[23] Fawzi, NL, Doucleff, M, Suh, JY, Clore, GM (2010) Mechanistic details of a protein-protein association pathway revealed by paramagnetic relaxation enhancement titration measurements. Proc Natl Acad Sci U S A 107: 1379–1384.

[24] Tang, C, Iwahara, J, Clore, GM (2006) Visualization of transient encounter complexes in protein-protein association. Nature 444: 383–386.

[25] Schilder, J, Ubbink, M (2013) Formation of transient protein complexes. Curr Opin Struct Biol 23: 911–918.

[26] Anthis, NJ, Clore, GM (2015) Visualizing transient dark states by NMR spectroscopy. Q Rev Biophys 48: 35–116.

[27] Dong, X, Qin, LY, Gong, Z, Qin, S, Zhou, HX, Tang, C (2022) Preferential interactions of a crowder protein with the specific binding site of a native protein complex. J Phys Chem Lett 13: 792–800.

[28] La, D, Kong, M, Hoffman, W, Choi, YI, Kihara, D (2013) Predicting permanent and transient protein-protein interfaces. Proteins 81: 805–818.

[29] Shi, Y, Pellarin, R, Fridy, PC, Fernandez-Martinez, J, Thompson, MK, Li, Y, Wang, QJ, Sali, A, Rout, MP, Chait, BT (2015) A strategy for dissecting the architectures of native macromolecular assemblies. Nat Methods 12: 1135–1138.

[30] Kastner, B, Fischer, N, Golas, MM, Sander, B, Dube, P, Boehringer, D, Hartmuth, K, Deckert, J, Hauer, F, Wolf, E, Uchtenhagen, H, Urlaub, H, Herzog, F, Peters, JM, Poerschke, D, Luhrmann, R, Stark, H (2008) GraFix: sample preparation for single-particle electron cryomicroscopy. Nat Methods 5: 53–55.

[31] Belsom, A, Rappsilber, J (2021) Anatomy of a crosslinker. Curr Opin Chem Biol 60: 39–46.

[32] Hofmann, T, Fischer, AW, Meiler, J, Kalkhof, S (2015) Protein structure prediction guided by crosslinking restraints--a systematic evaluation of the impact of the crosslinking spacer length. Methods 89: 79–90.

[33] Yu, C, Novitsky, EJ, Cheng, NW, Rychnovsky, SD, Huang, L (2020) Exploring spacer arm structures for designs of asymmetric sulfoxide-containing MS-cleavable cross-Linkers. Anal Chem 92: 6026–6033.

[34] Ding, YH, Fan, SB, Li, S, Feng, BY, Gao, N, Ye, K, He, SM, Dong, MQ (2016) Increasing the depth of mass-spectrometry-based structural analysis of protein complexes through the use of multiple cross-Linkers. Anal Chem 88: 4461–4469.

[35] Gong, Z, Ye, SX, Nie, ZF, Tang, C (2020) The conformational preference of chemical cross-linkers determines the cross-linking probability of reactive protein residues. J Phys Chem B 124: 4446–4453.

[36] Yang, B, Wu, H, Schnier, PD, Liu, Y, Liu, J, Wang, N, DeGrado, WF, Wang, L (2018) Proximity-enhanced SuFEx chemical cross-linker for specific and multitargeting cross-linking mass spectrometry. Proc Natl Acad Sci U S A 115: 11162–11167.

[37] Ziemianowicz, DS, Ng, D, Schryvers, AB, Schriemer, DC (2019) Photo-cross-linking mass spectrometry and integrative modeling enables rapid screening of antigen interactions involving bacterial transferrin receptors. J Proteome Res 18: 934–946.

[38] Wang, JH, Tang, YL, Gong, Z, Jain, R, Xiao, F, Zhou, Y, Tan, D, Li, Q, Huang, N, Liu, SQ, Ye, K, Tang, C, Dong, MQ, Lei, X (2022) Characterization of protein unfolding by fast cross-linking mass spectrometry using di-ortho-phthalaldehyde cross-linkers. Nat Commun 13: 1468.

[39] Kotrba, P, Inui, M, Yukawa, H (2001) Bacterial phosphotransferase system (PTS) in carbohydrate uptake and control of carbon metabolism. J Biosci Bioeng 92: 502–517.

[40] Deutscher, J, Francke, C, Postma, PW (2006) How phosphotransferase system-related protein phosphorylation regulates carbohydrate metabolism in bacteria. Microbiol Mol Biol Rev 70: 939–1031.

[41] Garrett, DS, Seok, YJ, Peterkofsky, A, Gronenborn, AM, Clore, GM (1999) Solution structure of the 40,000 Mr phosphoryl transfer complex between the N-terminal domain of enzyme I and HPr. Nat Struct Biol 6: 166–173.

[42] Garrett, DS, Seok, YJ, Peterkofsky, A, Clore, GM, Gronenborn, AM (1997) Identification by NMR of the binding surface for the histidine-containing phosphocarrier protein HPr on the N-terminal domain of enzyme I of the Escherichia coli phosphotransferase system. Biochemistry 36: 4393–4398.

[43] Cai, M, Williams, DC, Jr., Wang, G, Lee, BR, Peterkofsky, A, Clore, GM (2003) Solution structure of the phosphoryl transfer complex between the signal-transducing protein IIAGlucose and the cytoplasmic domain of the glucose transporter IICBGlucose of the Escherichia coli glucose phosphotransferase system. J Biol Chem 278: 25191–25206.

[44] Suh, JY, Tang, C, Clore, GM (2007) Role of electrostatic interactions in transient encounter complexes in protein-protein association investigated by paramagnetic relaxation enhancement. J Am Chem Soc 129: 12954–12955.

[45] Reizer, J, Sutrina, SL, Wu, LF, Deutscher, J, Reddy, P, Saier, MH, Jr. (1992) Functional interactions between proteins of the phosphoenolpyruvate:sugar phosphotransferase systems of Bacillus subtilis and Escherichia coli. J Biol Chem 267: 9158–9169.

[46] Chen, ZL, Meng, JM, Cao, Y, Yin, JL, Fang, RQ, Fan, SB, Liu, C, Zeng, WF, Ding, YH, Tan, D, Wu, L, Zhou, WJ, Chi, H, Sun, RX, Dong, MQ, He, SM (2019) A high-speed search engine pLink 2 with systematic evaluation for proteome-scale identification of cross-linked peptides. Nat Commun 10: 3404.

[47] Gong, Z, Ding, YH, Dong, X, Liu, N, Zhang, EE, Dong, MQ, Tang, C (2015) Visualizing the ensemble structures of protein complexes using chemical cross-linking coupled with mass spectrometry. Biophys Rep 1: 127–138.

[48] Matthew Allen Bullock, J, Schwab, J, Thalassinos, K, Topf, M (2016) The importance of non-accessible crosslinks and solvent accessible surface distance in modeling proteins with restraints from crosslinking mass spectrometry. Mol Cell Proteomics 15: 2491–2500.

[49] Combe, CW, Fischer, L, Rappsilber, J (2015) xiNET: cross-link network maps with residue resolution. Mol Cell Proteomics 14: 1137–1147.

